# A diet of oxidative stress-adapted bacteria improves stress resistance and lifespan in *C. elegans* via p38-MAPK

**DOI:** 10.1101/2023.10.06.561262

**Authors:** Ajay Bhat, Rebecca L. Cox, Brice Graham Hendrickson, Nupur K. Das, Emily Wang, Yatrik M. Shah, Scott F. Leiser

## Abstract

Organisms across taxa are exposed to stresses such as variable temperature, redox imbalance, and xenobiotics. Successfully responding to stress and restoring homeostasis is crucial for viability of the organism. During aging, the ability to effectively respond to stress declines, contributing to development of disease. In many multicellular animals, aging also coincides with changes in the microbiome that can contribute to disease-states. Because animals and their microbiota coexist in the same broad environment, they each must adapt to similar stresses. However, the short generation time of microbes leads to faster evolution, allowing the possibility that microbial stress adaptation may influence host physiology. Here we leverage a simplified model involving the nematode *C. elegans* and its bacterial diet. Our work highlights how bacterial adaptation to oxidative stress impacts the host’s lifespan and response to stress. Intriguingly, our findings reveal that worms fed with bacteria adapted to withstand oxidative stress exhibit enhanced stress resistance and an extended lifespan. Through whole genome sequencing, genetic assays, and metabolic analysis, this study underscores the pivotal role of the bacterial iron-sulfur pathway in governing host stress resistance and lifespan. We further find that iron in the stress-evolved bacteria boost the worm’s stress resistance and lifespan through activation of the mitogen-activated protein kinase (MAPK) pathway. In conclusion, this study provides evidence that understanding the evolutionary path of microbial adaptation during stress could be leveraged to slow aging and mitigate age-related decline in health.

## Introduction

Organisms spend their entire lives responding and adapting to environmental and systemic stressors such as oxidative stress, protein misfolding and nutritional imbalances. These responses are often carried out by the activation of varied stress response pathways^1^. The ability of an organism to counter such insults declines with age, leading to the accumulation of damage and contributing to systemic dysfunction and the progression of age-related disorders^1–3^. A notable example of such damage is the accumulation of reactive oxygen species (ROS) with age^4^. ROS are generated through aerobic metabolism and various environmental stressors that can damage cellular components including lipids, proteins and nucleic acids^5^. To combat ROS and maintain homeostasis, cells activate a set of signaling pathways known as the oxidative stress response^6^. When these responses fail, ROS accumulates in cells and is often associated with aging and the development of age-related diseases including diabetes, cardiovascular disease, cancer, Alzheimer’s disease, and Parkinson’s disease^4^.

Adding to the complexity of stress response, organisms do not live in isolation but are surrounded by and in many cases host trillions of microorganisms. The collection of microbes living on or inside other organisms, collectively called the microbiota, are also capable of sensing and responding to stress. The microbiota supplies essential metabolites, can aid in the digestion of complex macronutrients, and influences host metabolism, susceptibility to infection, and response to pharmaceuticals^7–9^. Emerging evidence suggests that microbes affect changes in host metabolism, stress response, and lifespan through the metabolites they secrete^10^. Additionally, the composition of the microbiota changes throughout the life of an organism and can be reflective of various disease states and overall health of the host^11,12^. Due to their short generation time and large population, microbes can evolve to resist stress much faster than their hosts^13,14^. This raises the question of whether bacterial adaptation to stress could result in physiological changes that modulate their hosts’ lifespan and susceptibility to that stress.

Iron is an essential nutrient that influences host-microbe interactions including microbial growth, host immunity function, and a range of biochemical processes^15^. Iron plays a critical role in several cellular processes, including oxygen transport, energy metabolism, transcription regulation, and DNA synthesis^16^. However, at high concentrations iron is toxic and acts a catalyst for the synthesis of ROS, leading to tight regulation of iron levels at both the cellular and the systemic levels^17^. During host-microbe interactions microbes utilize adaptive mechanisms to maintain homeostasis when iron levels fluctuate^18–20^. One of the major regulators of iron homeostasis in bacteria is the iron-sulfur (Fe-S) regulator *iscR*, which is a global transcriptional regulator of Fe-S biogenesis^21^. Interestingly, bacterial mutants adapted to the inflamed gut of aged mice frequently harbor *iscR* gene mutations^22^. Moreover, the fitness of these bacterial mutants is influenced by iron availability in the gut, hinting towards the importance of bacterial iron and the *iscR* gene in modulating host-microbe interactions^20^. However, the mechanism through which bacterial iron or iron-sulfur biogenesis regulates host stress susceptibility and lifespan remains unknown.

Previous work has shown that increased iron uptake in nematodes is correlated with an increased immune response^23^. The mitogen-activated protein kinase p38-MAPK, involved in the innate immunity pathway in *C. elegans*, is an evolutionarily conserved pathway that is activated by diverse environmental stresses, including xenobiotic and oxidative stress^24^. Much research has focused on the oxidative stress response and modulating MAPK targets to increase the host antioxidant capacity as a potential therapeutic for aging-related diseases from cancer to neurodegeneration^24,25^. The *C. elegans* p38-MAPK homolog, PMK-1, plays an important role in survival during exposure to pathogenic bacteria and oxidative stress and regulates the induction of genes involved in maintaining redox homeostasis^25,26^. Given the role of worm stress pathways in response to xenobiotics, it may be possible for bacterial adaptations to induce broader signaling changes in the host including those governed by p38-MAPK.

Due to the complexity of the mammalian microbiome, assigning causality to microbial genes on host stress response and longevity is both costly and technically challenging^27^. The nematode *Caenorhabditis elegans,* which has a short lifespan and a well-defined microbiota, can be a useful model for the actions of a simplified microbiome. Typically, *C. elegans* is grown in laboratory conditions with one strain of *Escherichia coli*, which colonizes the intestinal lumen and forms the gut microbiota^28^. With the aid of this simplified host-microbe model, we can explore various roles microbes play in regulating the stress response and longevity of the host^29^. Metabolites provided by different bacterial strains can affect the longevity of the worms, including Vitamin B12, nitric oxide, and colanic acid. Many of these pro-longevity metabolites modulate lifespan by regulating the worm stress response^30,31,32^. However, it is still unclear whether changes in the microbial stress response can modulate host stress signaling to regulate health and lifespan.

In this study, we test whether modifying the microbiome can directly affect host longevity and susceptibility to stress. We use adaptive laboratory evolution to genetically modify OP50, a strain of *E. coli* commonly used as a food source for *C. elegans*, to resist oxidative stress. We demonstrate that worms grown on these bacteria are also resistant to oxidative stress and have increased lifespan. Using whole genome sequencing, we identify a mutation in the iron sulfur cluster regulator (*iscR*) gene in our lab-evolved bacteria that contributes to increased host stress resistance and demonstrates the role of the bacterial Fe-S pathway in regulating worm lifespan and stress tolerance. Additionally, we find that stress-evolved bacteria mediate the worm’s stress resistance via the PMK-1/p38-MAPK signaling pathway. Our results support a model where adaptations in stress resistance of the microbiota have the potential to induce hormetic stress responses in the host, improving the trajectory of host health.

## Results

### Paraquat-evolved bacteria increases the stress tolerance and lifespan of worms

To test if bacteria adapted to resist ROS can alter the stress tolerance and lifespan of worms, we used adaptive laboratory evolution (ALE). We evolved OP50 bacteria, an *E. coli* strain routinely used as a *C. elegans* food source in laboratory conditions, in the presence of the ROS inducer, paraquat. We chose paraquat because of the association between aging and oxidative stress-mediated damage^4^, and because paraquat is the most consistent stress in our previous cell culture and worm studies^33,34^. During ALE, bacteria are continuously grown under stress conditions to allow for the selection of the improved stress resistance phenotype. Genetic diversity can increase the probability of beneficial genotypes and phenotypes. Various studies have used techniques like genome shuffling, chemical mutagenesis, and multiplexed automated genome engineering to generate genomic diversity during the ALE process^35^. We used ethylmethanesulfonate (EMS) to generate random mutations, followed by ALE to select mutants resistant to paraquat. Using this combined approach, we evolved OP50 for roughly 500 generations in the presence of 2.5 mM paraquat to generate a <u>P</u>ara<u>q</u>uat <u>E</u>volved OP50 strain (OP50^PQE^) **(Fig 1A).** Using growth kinetics and dose-response we find that OP50^PQE^ bacteria grow better in the presence of paraquat **(Fig. S1A)** and resist a higher range of paraquat concentrations **(Fig. 1B).**

**Fig 1.**
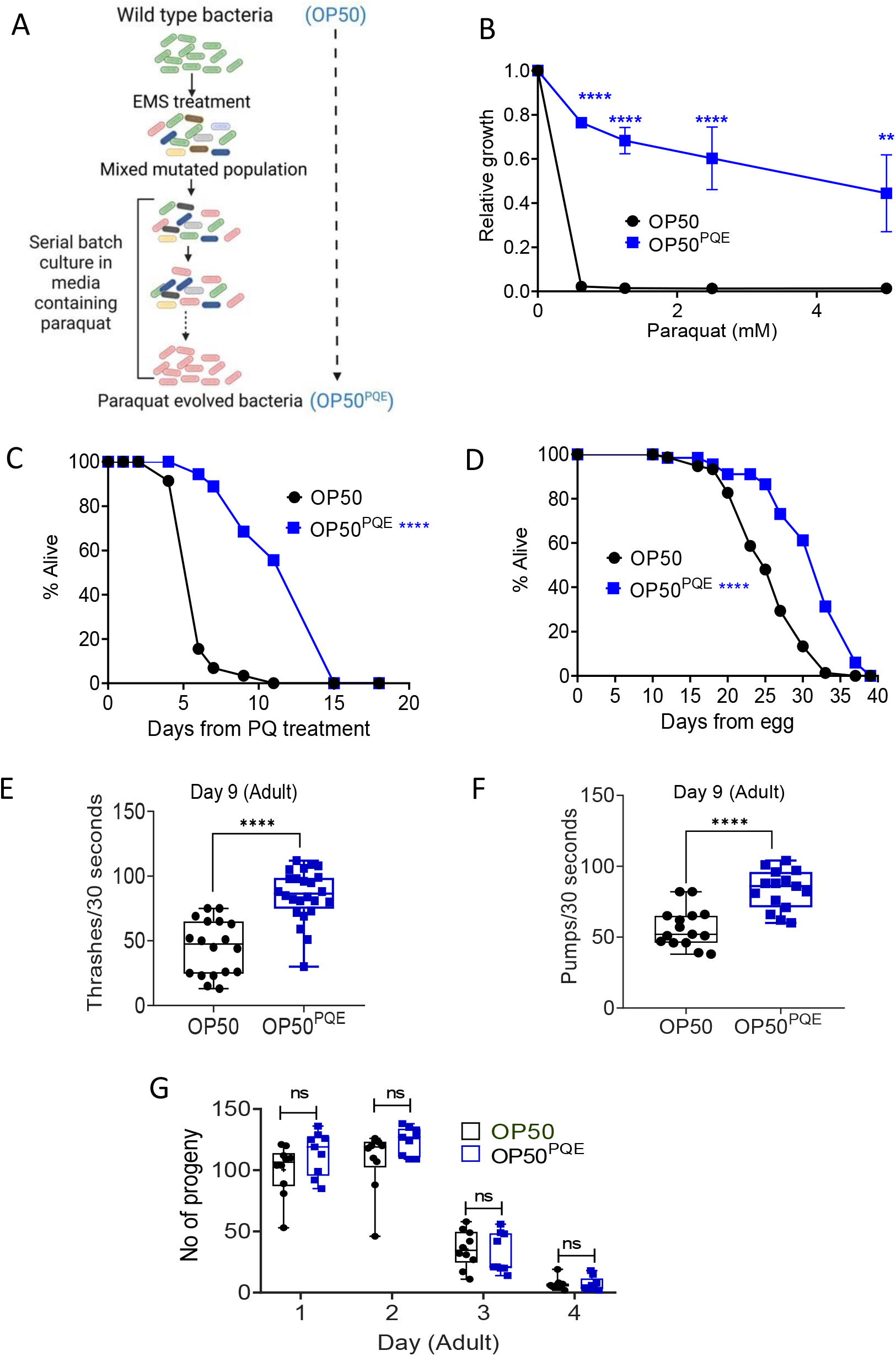
Bacteria adapted to resist paraquat increase worm oxidative stress resistance, health and lifespan. A. Schematic of guided evolution strategy. OP50 *E. coli* was mutagenized with EMS followed by laboratory adaptive evolution by growing in increasing concentrations of paraquat until resistance was stable. B. Dose-dependent effect on growth of different concentrations of paraquat on the growth of OP50^PQE^ and OP50. Error bar represents mean +/- SEM. ** and **** represent p-value < 0.01 and <0.0001, respectively, using two-way ANOVA. n = 3 independent replicates. C. Survival curve of worms fed OP50 or OP50^PQE^ in the presence of 2.5 mM paraquat. Worms fed either bacteria were grown until L4 stage and transferred to FUdR plates containing 2.5 mM paraquat. **** represents p-value < 0.0001 calculated using log-rank test. D. Lifespan assay comparing the survival of worms fed OP50^PQE^ or OP50 bacteria. **** represents p-value < 0.0001 calculated using log-rank test. E. Thrashing assay of day 9 adult worms fed OP50^PQE^ or OP50 bacteria. Box plot represents the number of thrashes per 30 seconds. **** represents p-value < 0.0001 calculated using unpaired t-test. n = 18-24 independent worms. F. Pharyngeal pumping of worms fed either OP50^PQE^ or OP50 bacteria. **** represents p-value < 0.0001 calculated using unpaired t-test. n = 15 independent worms. G. Reproductive capacity of worms fed OP50^PQE^ or OP50 bacteria. Number of progeny was calculated from individual worms across different days of adulthood. ns = not-significant, p-value calculated using two-way ANOVA Worm stress assay and lifespan statistics are in Supplementary data 1.

We next tested whether growing worms on the OP50^PQE^ strain impacts their stress tolerance and lifespan. Interestingly, we find that worms grown on OP50^PQE^ are significantly more resistant to paraquat than the worms grown on the wild type OP50 **(Fig. 1C).** This result suggests that bacteria that have evolved to resist ROS can improve the stress tolerance of worms. Increased stress resistance often correlates with increased lifespan ^36^, so we also tested whether the OP50^PQE^ bacteria affected worm longevity. We measured the lifespan of worms grown on OP50 and OP50^PQE^ bacteria and find that worms grown on OP50^PQE^ live longer **(Fig. 1D)**. We confirmed that this increase in lifespan correlates with improved healthspan, measured by the increased thrashing and pumping rates in middle-aged worms fed OP50^PQE^ (**Fig. 1E-F**). Lifespan extension is often associated with decreased fecundity, leading to smaller brood size ^37^. However, we find that the overall brood size (**Fig. S1B**) and rate of egg-laying does not change between the two bacterial conditions (**Fig. 1G**). These results suggest that modulating microbial stress tolerance can positively influence the lifespan and healthspan of its host.

### Bacterial Fe-S cluster biogenesis pathway regulates worms stress tolerance and lifespan

To identify the genetic changes responsible for paraquat tolerance, we sequenced the genome of the paraquat-evolved bacteria (OP50^PQE^). We find that OP50^PQE^ has 36 mutations across different genes in comparison to the wild-type strain (OP50) **(Supplementary data 2**). A majority of these mutations are missense mutations (**Fig. S2A**). Three of these mutations are associated with genes involved in stress response (**Fig. S2B**), including *iscR* (iron-sulfur cluster transcriptional regulator), *rpoH* (RNA polymerase sigma factor), and *grpE* (nucleotide exchange factor). The *iscR* mutation is of particular interest, since a separate study reported a mutation in the same gene while evolving *E. coli* bacteria with paraquat^38^. This underlines the importance of the *iscR* gene in the context of paraquat-mediated adaptive evolution. The IscR protein regulates the expression of genes that encode the Fe-S cluster assembly proteins, IscS, IscU, and IscA. IscS is a cysteine desulfurase, which donates sulfur to the Fe-S cluster assembly by converting cysteine to alanine (**Fig. 2A**). The scaffold protein IscU uses free iron or iron transferred from IscA to assemble the Fe-S cluster. The complete cluster is then transferred to IscA, which acts as a carrier and transfers the Fe-S cluster to apo-enzyme substrates^39^. Based on our data showing an *iscR* mutation in OP50^PQE^, we examined the effect of the deletion of the ISC genes on bacterial sensitivity to paraquat, using bacterial strains from the *E. coli* K12 Keio knock-out library. We find that loss of all ISC genes except *iscR* render bacteria sensitive to paraquat, while *iscR* deletion enhances paraquat resistance (**Fig. 2B**).

**Fig 2.**
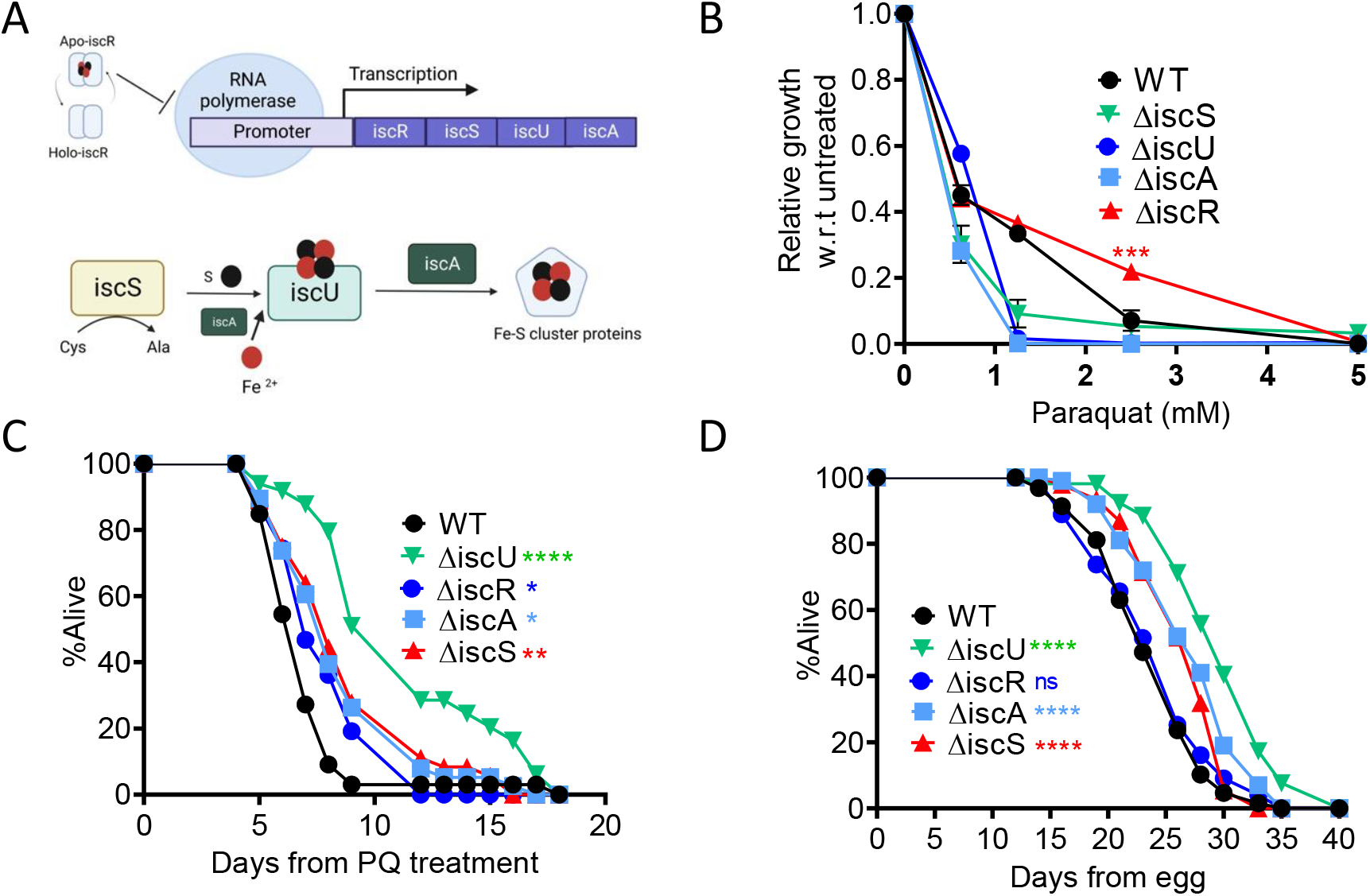
Bacterial Fe-S cluster biogenesis pathway regulate worms stress tolerance and lifespan. A. Schematic of ISC operon and Fe-S biogenesis B. Effect of different concentration of paraquat on the growth of isc KO strains. *** represents p-value < 0.001 comparing growth of WT vs ΔiscR bacteria using 2-way ANOVA. p-value for other strains (ΔiscU, ΔiscA, and ΔiscS) is <0.0001 across all the concentrations of paraquat except for 5 mM (non-significant). Data are represented as mean +/- SEM. n = 3 independent biological replicates. C. Survival curve of worms fed WT or different isc bacterial mutants in presence of paraquat. Worms fed either of the bacteria were grown till L4 stage and transferred to FUdR plates containing 5 mM paraquat. *, **, and **** represent p-value < 0.05, < 0.01 and < 0.0001 calculated using log-rank test. D. Lifespan assay comparing the survival of worms fed wild type or isc mutants. **** represents p-value < 0.0001 calculated using log-rank test. Worm stress assay and lifespan statistics are in Supplementary data 1.

We noted that the *iscR* deletion in the K12 strain does not increase paraquat resistance as dramatically as the point mutation observed in OP50^PQE^. This result is plausibly due to differences in the background bacterial strains, as results from **Fig. 1B** and **Fig. 2B** suggest different basal levels of resistance to paraquat between the strains. We thus asked if there is a significant difference between OP50 and K12 bacterial paraquat resistance and whether this translates to differences in worm resistance. Indeed, we find K12 control strains are more resistant to paraquat than OP50 (**Fig. S2C**) and that this enhanced resistance is also apparent in worms grown on K12 (**Fig.S2D**). These data further support the hypothesis that bacterial paraquat resistance can improve worm paraquat resistance.

Given our observation that iron-sulfur cluster operon mutations affect bacterial stress resistance (**Fig 2B**), we next asked how these bacterial mutations affect worms. Interestingly, worms grown on ISC KO strains, Δ*iscR*, Δ*iscA*, Δ*iscU* and Δ*iscS* are each resistant to paraquat, with Δ*iscU* showing the strongest effect (**Fig. 2C**). Measuring lifespan, we find that all ISC mutants except *iscR* increase the longevity of worms **(Fig 2D).** These results indicate that perturbations of any one of the ISC assembly genes greatly affects the physiological response of bacteria to paraquat exposure in a manner that causes increased stress resistance and lifespan of the host. However, while the *iscR* mutation in OP50^PQE^ bacteria leads to improved stress tolerance and extended lifespan in worms, the *iscR* deletion in K12 bacteria only increased worm resistance. This discrepancy could be due to differences between the OP50^PQE^ and Δ*iscR* strains, or it could result from differing effects caused by *iscR* gene deletion compared to a single amino acid variation within the protein. While the assembly genes and regulators can have differing effects on lifespan, these results highlight that bacterial iron-sulfur biogenesis influences worm stress tolerance and lifespan.

### Higher levels of iron in OP50^PQE^ improve bacterial growth in paraquat stress conditions

Given the mutation in the bacterial iron sulfur cluster operon, we hypothesized that iron levels may be important for stress phenotypes in either the bacteria, the worms, or both. To address this, we first asked whether the basal level of iron in the bacterial strain was changed. Using inductively coupled plasma mass spectrometry (ICP-MS), we found a remarkable 50% increase in the iron content of OP50^PQE^ compared to the wild type OP50. This increased level was sustained after the addition of 5 mM paraquat to the growth media (**Fig 3A**). To test if the increased iron levels were necessary for the higher resistance to paraquat of the adapted bacteria, we performed an overnight growth assay in which 2,2’-bipyrydyl, an iron chelator, was added to the growth media (**Fig 3B**). The resistance of OP50^PQE^ significantly decreased with the addition of 250 mM of the chelator, suggesting that high levels of iron in OP50^PQE^ are largely responsible for paraquat resistance.

**Fig 3.**
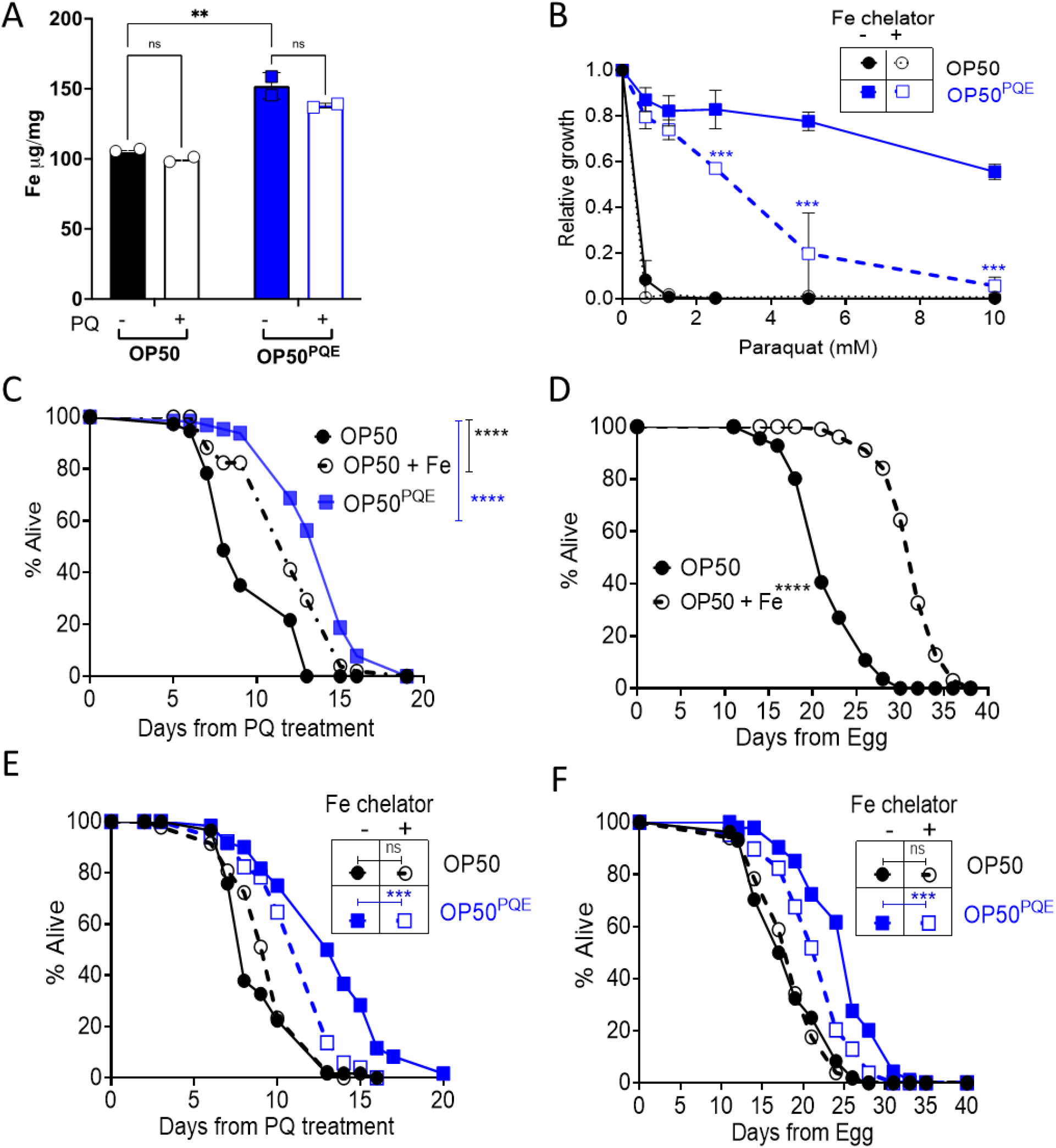
Bacterial iron levels modulate the worms’ paraquat tolerance. A. Levels of iron in OP50 and OP50^PQE^ bacteria with and without paraquat treatment measured by ICP-MS. Overnight bacterial culture was incubated with or without 2.5 mM of paraquat for one hour before being processed for Fe measurement. Data represent mean +/- SEM, and p-value is calculated using one-way ANOVA. B. Dose-dependent effect of different concentrations of paraquat on the growth of OP50 and OP50^PQE^ bacteria in the presence and absence of 250 µM 2,2-bipyridyl (Fe chelator). Data represent mean normalized growth with respect to the untreated control condition. The error bar indicates SEM, and p-value is calculated using two-way ANOVA. C. Paraquat survival assay of worms fed OP50 or OP50^PQE^ in the presence of iron. Worms fed OP50 or OP50^PQE^ were transferred at L4 stage to FUdR plates seeded with either of the bacteria mixed with or without 5 mM Fe. p-value was calculated using log-rank test. D. Effect of Fe on the lifespan of worms fed OP50 bacteria. Worms fed OP50 were transferred as day 1 adults to FUdR plates seeded with OP50 mixed with or without 5 mM Fe. p-value was calculated using log-rank test. E. Paraquat survival assay of worms fed OP50 or OP50^PQE^ treated with 2,2-bipyridyl Fe chelator. Worms fed OP50 or OP50^PQE^ were transferred at L4 stage to FUdR plates containing 2.5 mM paraquat seeded with either of the bacteria mixed with or without Fe chelator. p-value was calculated using log-rank test. F. Lifespan of worms fed OP50 or OP50^PQE^ treated with 2,2-bipyridyl Fe chelator. Worms fed OP50 or OP50^PQE^ were transferred as day 1 adults to FUdR plates containing 2.5 mM paraquat seeded with either of the bacteria mixed with or without Fe chelator. p-value was calculated using log-rank test. **, *** and **** represents p-value < 0.01, < 0.001 and < 0.0001, respectively. Worm stress assay and lifespan statistics are in Supplementary data 1.

### Higher iron levels are required for the ability of OP50^PQE^ to improve worm paraquat stress tolerance

We next asked whether supplementing iron in the OP50 food source would be sufficient to phenocopy the OP50^PQE^-mediated worm stress resistance to paraquat. Interestingly, of the conditions tested, only the addition of 5 mM iron to OP50 consistently increased worm stress resistance to levels comparable to OP50^PQE^-fed worms (**Fig 3C**). All concentrations of iron tested significantly decreased the resistance of OP50^PQE^-fed worms, but lower concentrations of iron did not significantly change the worms’ paraquat tolerance when fed OP50 (**Fig. S3A, B**). Iron supplementation of OP50 also mimics the increased lifespan of worms fed OP50^PQE^ alone (**Fig. 3D**). Conversely, mixing an iron chelator with seeded bacteria had no significant effect on the resistance of worms fed OP50, but significantly decreased the resistance and lifespan of worms fed OP50^PQE^ (**Fig. 3E, 3F**). This indicates a somewhat narrow range of beneficial iron concentrations for paraquat tolerance.

### pmk-1/p38-MAPK is required for OP50^PQE^-induced paraquat stress resistance

Having established that laboratory-adapted bacteria can induce stress resistance and longevity in *C. elegans,* dependent on iron, we were left with the question of mechanism in the worms. Given that paraquat is known to cause mitochondrial stress, and iron is an important component in many mitochondrial proteins, we asked if the mitochondrial stress response could be altered in worms fed OP50^PQE^. We examined expression of *hsp-6*, a reporter for mitochondrial stress that is induced by paraquat exposure, using a fluorescent reporter strain. Our results show that worms grown on OP50 or OP50^PQE^ have similar levels of *hsp-6* induction both at a basal level (**Fig. S4A**) and when exposed to paraquat (**Fig. S4B**).

Given this result, we wondered what other pathways might be involved. We performed RNA-seq of worms fed wild type vs paraquat adapted OP50 in the presence and absence of paraquat. We found that genes involved in the innate immune response were highly enriched in worms fed with either of the two bacteria and exposed to paraquat (**Fig. S4C, Supplementary data 3,4 and 5**). Interestingly, we find that genes involved in the innate immune response are upregulated in worms fed OP50^PQE^ even in the absence of paraquat (**Fig 4A, Supplementary data 6 and 7**). Using WormExp enrichment tool^40^, we find that many of the differentially upregulated genes are downstream of the *C. elegans* p38-MAPK homolog, PMK-1 (**Fig. S4D)**. Using a reporter strain of *irg-5,* a target of PMK-1 involved in the worm innate immune response, we confirmed that worms grown on OP50^PQE^ have significantly higher expression of this gene (**Fig 4B, C**). We find that PMK-1 is required to confer stress resistance, as the *pmk-1* mutant has similar paraquat tolerance when fed either OP50 or OP50^PQE^ (**Fig 4D**). We also tested the role of *sek-1*, which is upstream of PMK-1, and find that *sek-1* mutants also do not benefit from OP50^PQE^ when faced with paraquat stress (**Fig. S4E**).

**Fig 4.**
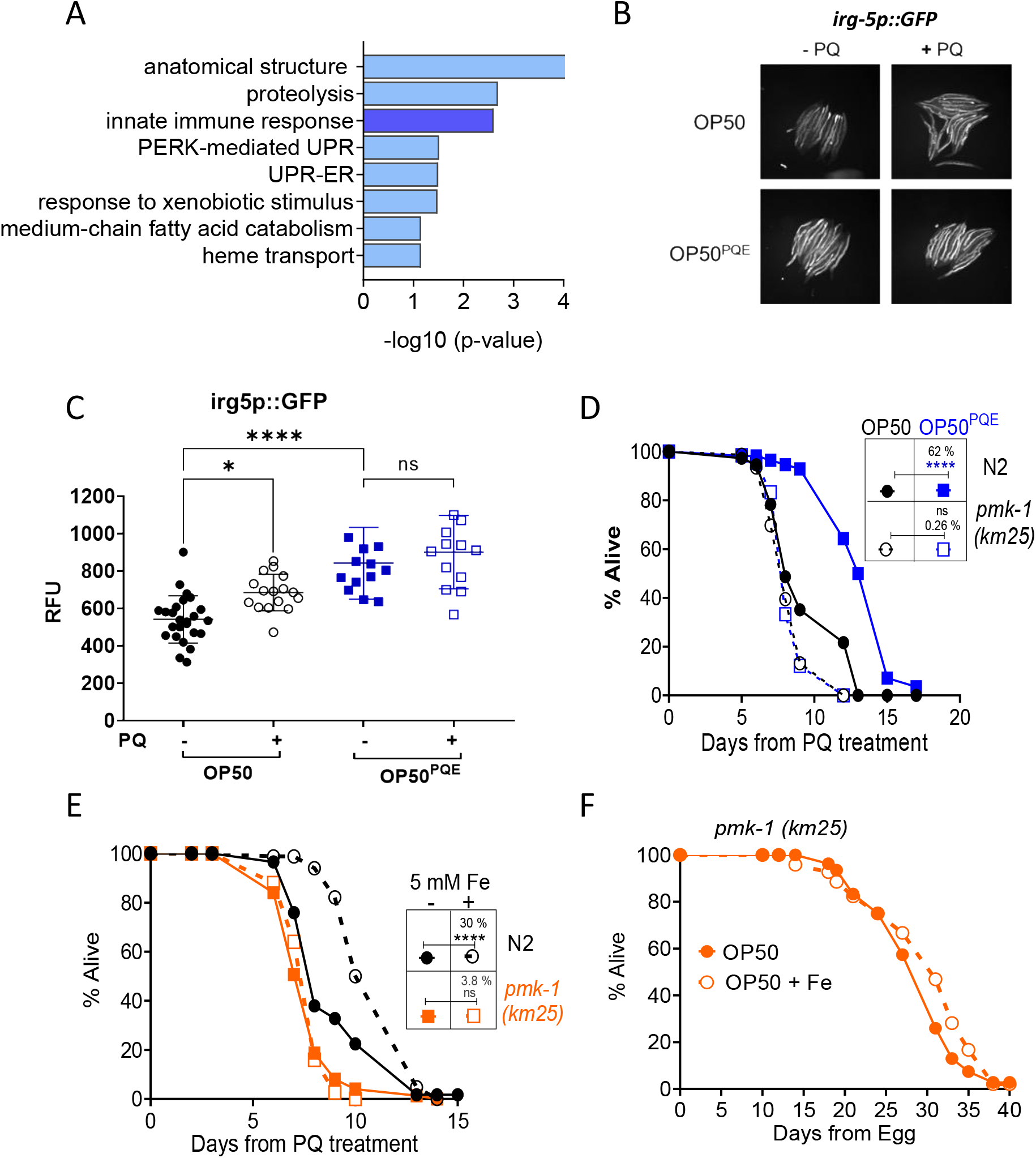
The effect of OP50^PQE^ bacteria on worms requires *pmk-1*. A. Biological terms enriched in worms grown on OP50^PQE^ bacteria vs OP50 bacteria. Analysis was generated using the DAVID bioinformatics resource (LHRI). B. Representative images of *irg-5p::GFP* reporter worms fed OP50^PQE^ bacteria vs OP50 bacteria on control and paraquat stress exposure conditions. C. Quantification of *irg-5p*::GFP reporter strain fluorescence. Error bars represent mean +/- SEM. * represents p < 0.05, **** < 0.0001 using an unpaired t-test. n = 15-20 independent worms. D. Survival curve of wild type (N2) or *pmk-1 (km25)* worms fed OP50 or OP50^PQE^ in the presence of 2.5 mM paraquat. Worms were transferred from NGM to FUdR and paraquat containing plates at L4. **** represent p < 0.0001 for a log-rank test. E. Survival curve of N2 and *pmk-1 (km25)* worms fed OP50 with and without iron supplementation exposed to paraquat. Worms were grown from egg on each respective bacteria and transferred to FUdR and 2.5 mM paraquat plates at L4. **** represent p < 0.0001 calculated using log-rank test. F. Lifespan curve of *pmk-1 (km25)* worms fed OP50 alone or mixed with 5 mM iron (OP50 + Fe). Worm stress assay and lifespan statistics are in Supplementary data 1.

Because high iron levels in OP50^PQE^ are required for increased stress tolerance in worms **(Fig. 3D)**, we investigated whether *pmk-1* plays a role in Fe-mediated stress resistance. We find that *pmk-1* is also required for iron supplementation to improve worm stress tolerance on OP50. This further supports the idea that increased bacterial iron levels are responsible for a significant portion of the beneficial effect of OP50^PQE^ for paraquat stress resistance (**Fig. 4E**). We also find that *pmk-1* is required for the OP50^PQE^ mediated lifespan extension (**Fig. 4F**).

## Discussion

Microbes live in proximity to their hosts and are often subjected to similar stress conditions. This raises the fundamental question of whether microbial adaptation during stress could affect host physiology. In this study we aimed to unravel how bacteria adapted to oxidative stress impact the stress tolerance and lifespan in host *C. elegans* that consume these bacteria. We discovered that worms fed evolved bacteria (OP50^PQE^) have enhanced stress tolerance and live longer. Our data indicate that these phenotypes are likely due in large part to a mutation in the bacterial iron-sulfur cluster pathway, which other bacterial evolution and mammalian microbiome studies have established as an important locus for evolutionary adaptation to oxidative stress^22,38^. Importantly, this study establishes a link between bacterial iron metabolism and worms’ p38 MAPK (PMK-1) signaling pathway, which regulates stress resistance and lifespan. These findings support the hypothesis that microbial adaptation to stress conditions can more broadly influence host organism health and gene expression.

Our study finds that ROS-adapted bacteria (OP50^PQE^) have a mutation in the iron-sulfur cluster regulator gene (*iscR*). Other work has found mutations in the same gene in paraquat-guided evolution experiments, demonstrating its importance in ROS-mediated adaptive evolution^38^. Furthermore, this gene is highly mutated in the bacteria colonizing the guts of 18-month-old mice, suggesting that its role in regulating host-microbe interactions may be conserved across species^22^. We found that deleting any of the iron-sulfur assembly proteins in bacteria has a positive effect on the stress tolerance and lifespan of worms, with the maximum effects seen with deletion of the *iscU* gene. Consistent with these results, suppressing the expression of *iscu-1*, a worm ortholog of *iscU*, promotes stress tolerance and longevity in worms^41^. Taken together, these findings suggest that disturbing Fe-S cluster homeostasis in either bacteria or worms can influence stress responses and longevity. Further studies are required to determine whether bacterial Fe-S cluster biogenesis influences the Fe-S cluster pathway in worms to regulate longevity. Fe-S clusters play an important role in the electron transport chain and tricarboxylic acid cycle^42^, thus perturbing the expression of Fe-S cluster genes in bacteria or host may modulate the host mitochondrial metabolism to regulate aging. In mammals, mitochondria are believed to have descended from bacteria as per the endosymbiotic theory, and play a crucial role in the biogenesis of Fe-S sulfur clusters^43^. Further investigation is required to ascertain whether microbes utilize the Fe-S cluster pathway to communicate with host mitochondria during external stress or aging.

The results from our study also tie into the complex system of iron homeostasis in the host and microbiota. Microbes competes with the host for iron, and iron availability in the host is tightly regulated to prevent the colonization of pathogenic bacteria^44^. In turn, microbes have evolved mechanisms to acquire iron from the host, including the secretion of iron-chelating molecules like siderophores^45^. Evolutionary pressure prevents organisms from simply increasing their intake of iron as high levels create oxidative stress due to the Fenton reaction, which converts hydrogen peroxide into hydroxyl radicals, a highly reactive oxygen species^46,47^. At the same time, iron is a cofactor for antioxidant enzymes, and low levels of iron also cause oxidative stress, suggesting an optimal window of iron levels is critical to maintaining redox homeostasis^48^.

We find that bacteria adapted to resist ROS have a mutation in the *iscR* gene and have high levels of iron, suggesting that bacterial adaptation to high levels of ROS target iron metabolism to increase oxidative stress resistance. Interestingly, these bacterial adaptations help worms to live longer and better tolerate oxidative stress. It is possible that bacteria adapted to ROS are either assisting the worms in achieving optimum iron levels or have reached a threshold that causes hormetic stress, which in turn results in worm stress tolerance and longevity^49^.

Our study also demonstrates that OP50^PQE^ bacteria activate p38 MAPK (PMK-1) targets in *C. elegans,* and that this pathway is required for OP50^PQE^ to benefit stress tolerance and lifespan. PMK-1 plays a crucial role in host-microbe interactions by activating the host’s innate immune response against pathogenic microorganisms^26^. This raises the question of why ROS-evolved bacteria trigger the innate immune system. A recent study shows that in long-lived *C. elegans* mitochondrial mutants such as *nuo-6* and *isp-1*, innate immunity genes including PMK-1 targets are enriched^50^. Thus, it is possible that mitochondrial metabolism may be altered by high levels of iron in the OP50^PQE^ bacteria, resulting in PMK1 activation. A second possibility is that high iron levels in OP50^PQE^ bacteria increase their colonization of worm intestines, which is detected as stress by the host and activates the immune system. In either case, this activation of innate immunity may allow the worms to combat future ROS insults.

Our data support a model where bacterial adaptation under ROS can alter Fe-S cluster biogenesis, increasing iron levels in the bacteria (**Fig. 5**). Iron from the bacteria plausibly activates PMK-1 signaling in worms, resulting in increased stress tolerance and an extended lifespan. Our data suggest that this beneficial interaction is not exclusively attributable to the resistance of the bacteria to ROS, as mutant ISC bacteria that are sensitive to ROS may also aid the worms in resisting ROS and extending their lifespan. Thus, the beneficial effect of these microbes could be due in part to secondary changes induced by altered stress tolerance, such as increased iron levels in the case of OP50^PQE^ bacteria. However, these mutations sensitizing the bacteria to stress are less likely to appear in the wild due to selective pressure promoting mutations that increase fitness. This indicates that it may be worthwhile to broadly examine how perturbing these pathways may benefit the host despite potential negative effects on microbe fitness.

**Fig 5.**
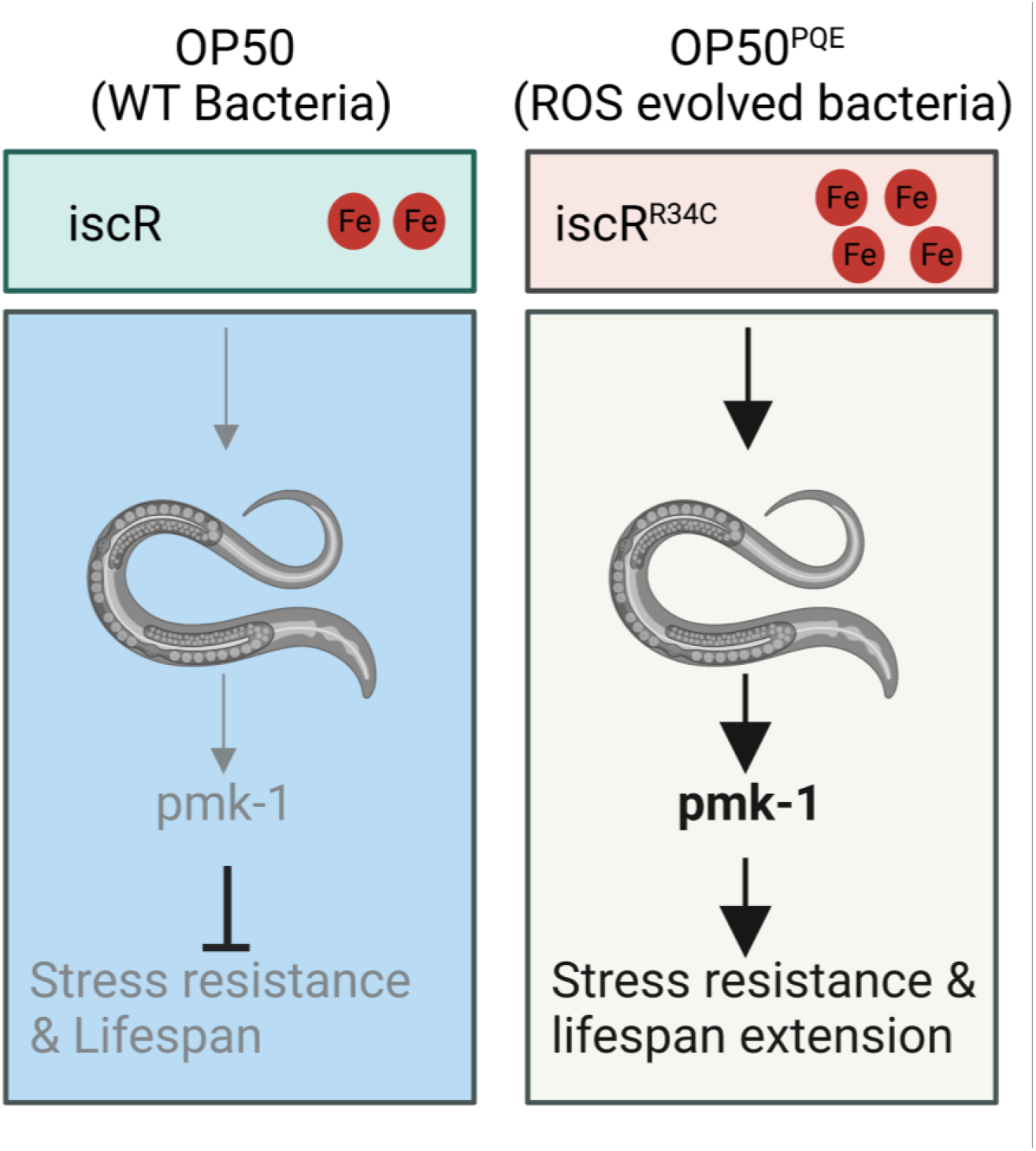
Proposed model: Mutation of the iscR gene in the evolved bacteria (OP50^PQE^) results in elevated iron levels compared to the wild type bacteria (OP50). Increased bacterial iron levels enhance stress tolerance and lifespan in worms via PMK-1 signaling.

Taken together, our study provides evidence that microbial stress signaling influences the host’s stress response and lifespan. By utilizing a simplified host-microbe model, we discovered that perturbing the Fe-S cluster biogenesis pathway in bacteria extends lifespan and enhances stress tolerance in *C. elegans* through the activation of the PMK-1 signaling pathway. Our study supports the notion that comprehending the evolutionary path of bacteria’s adaptation during stress can be leveraged to decelerate the aging process and alleviate age-related diseases. However, further research is required to determine if this interaction is specific to ROS and whether it can be translated to mammalian systems with complex microbiota.

## Materials and Methods

### Strains and growth conditions

*C. elegans* were maintained using standard methods as previously described^33,51^. Wild-type N2, SJ4100 *(zcIs13 [hsp-6p::GFP + lin-15(+)])*, AY101 *(acIs101 [irg5p::GFP + rol-6(su1006)]*, KU25 *(pmk-1 (km-25)),* KU4 *(sek-1 (km4))* were grown on NGM (nematode growth media) plates seeded with OP50 *E. coli*, OP50^PQE^ adapted *E. coli* or K12 Keio library strains to match the bacteria used in the intended experiments. OP50 bacteria and worm strains were obtained from CGC, and K12 Keio library strains were purchased from Horizon Discovery Ltd. All worm experiments were performed at 20°C. Some strains were provided by the CGC, which is funded by NIH Office of Research Infrastructure Programs (P40 OD010440).

### Bacterial growth conditions

Bacterial cultures were grown overnight in Luria broth from single colonies. Cultures for OP50 and K12-derived bacteria were shaken at 225 RPM at 37°C for 16 hours.

### Bacterial adaptive evolution

OP50 *E. coli* were adapted to grow in 2.5 mM paraquat (methyl viologen dichloride hydrate, Sigma Aldrich 856177) following a modified adaptive evolution protocol ^52,53^. Overnight grown OP50 were re-inoculated in fresh LB media (2 ml) and grown till it reached ∼ 0.3 O.D, and then the cells were washed twice with phosphate buffer saline (PBS, pH 7.5). Cells were then reconstituted in PBS (2 ml) and were treated with EMS and incubated at 37 °C for 45 minutes. Cells were then washed twice with PBS and allowed to recover in LB at 37 °C for 2-3 hours. Cultures were first exposed to 0.5 mM paraquat, and concentration was increased over subsequent generations and then finally sub-cultured in 2.5 mM paraquat for ∼500 generations. Diluted culture was then plated on LB agar plates to isolate single colonies.

### Stress resistance assay

To induce oxidative stress, we added 2.5 mM or 5 mM paraquat to FUdR plates not containing antibiotic (40690016, Bioworld). Equal O.D of bacteria were seeded and allowed to dry for 48 hours prior to transferring worms to the plates. Gravid adult worms were placed on NGM plates seeded with experimental bacteria for a timed egg-lay, and after 3 hours were removed and synchronized eggs were allowed to develop at 20°C. When the worms reach L4 stage they were transferred to paraquat-containing plates, with at least 2 plates per replicate and minimum 40 worms per plate. One additional transfer to paraquat plates was performed 24 hours later, at which point worms were counted and scored every day. Once worms did not respond to gentle nudging with a worm pick they were counted as dead. A similar method was used for paraquat stress assays with the addition of iron (Iron (III) sulphate hydrate, Thermo Fisher 033316.30) or iron chelators (2,2’-Bipyridyl, Sigma Aldrich D216305), in which the compound was mixed with bacteria prior to seeding. Iron and Iron chelator both were dissolved in water with the final stock of 500 mM and 17.5 mM, respectively.

### Bacterial stress resistance assays

The OD_600_ of bacterial overnight cultures was measured, and cultures were diluted to a O.D of 0.003 in LB media. 400 uL of these dilutions were added to 2 mL deep well plates, and paraquat was added at its highest concentration (5 mM) and serially diluted in each bacterial culture. Cultures were grown in deep well plates for 16-18 hours with a gradient of paraquat at which point samples were measured at 600 nm on a Biotek plate reader. To correct for background signal, the O.D from the blank well was subtracted from the O.D measured from the experimental wells. These values were then normalized by those obtained from their respective untreated samples.

### Lifespans

Worms were maintained on NGM plates seeded with the bacteria to be used in the lifespan assay for one generation. Gravid adults were placed on NGM seeded with the appropriate bacteria to lay eggs for 3 hours and then removed. Once this F1 generation reached the young adult/gravid adult stage, worms were transferred to plates containing FUdR, with 50 worms per plate and a minimum of 2 plates per condition in each replicate lifespan. 3 more transfers were performed at day 2, 5, and 7 of adulthood, and deaths were counted beginning at day 9 of adulthood and were counted and scored every other day. Once worms did not respond to gentle nudging with a worm pick they were counted as dead.

### Pumping and thrashing

Worms were prepared the same as for lifespans and transferred to FUdR plates at day 1 of adulthood. Worms were first transferred to an empty plate and then moved to a drop of M9 buffer and filmed for a total of 30 seconds. Each body bend was counted during a thirty second period of maximal movement. A minimum of twenty worms per condition were filmed per replicate. Pumping was measured on a Leica M205 C dissecting stereomicroscope at 100x magnification. Movement of the pharynx was counted for thirty seconds in a minimum of ten worms per replicate.

### Brood size assay

Gravid adult worms were placed on NGM plates seeded with control or experimental bacteria for a one hour timed-egg lay. Progeny were moved individually to 35 mm plates seeded with the same bacteria at L3/L4 stage. When worms began laying eggs they were transferred to fresh plates every 24 hours. Live progeny were counted after reaching L3 stage. A minimum of ten worms were grown on each bacterial strain per replicate, for a total of three replicates.

### RNA isolation and transcriptomic analysis

Approximately 500 adult worms per biological replicate were washed three times with M9 buffer, then frozen in liquid nitrogen and stored at -80 °C. During RNA extraction 500 ml of Trizol reagent was added to the frozen pellets, followed by three freeze-thaw cycles with liquid nitrogen and water bath at 42°C. Then 500 ul of ethanol was added to the tubes, and the total RNA was isolated using the Direct-Zol Miniprep Plus Kit (Cat#R2072). Purified RNA was sent to Novogene (Novogene Corporation Inc) for sequencing on the Illumina HWI-ST1276 instrument. The poly-T oligo-attached magnetic beads were used to purify messenger RNA from total RNA. Following fragmentation, random hexamer primers were used to synthesize first strand cDNA, followed by second strand cDNA. Library construction included end repair, A-tailing, adapter ligation, size selection, amplification, and purification. The library was then sequenced on an Illumina device using paired-end sequencing. Gene count data obtained from RNA-seq analysis were passed into the DESeq2 analysis package in R. The results output by DESeq2 were sorted by p-value to obtain a list of genes most likely to be differentially expressed due to stress-evolved diet.

### Bacterial genome sequencing and analysis

Six individual clones of OP50^PQE^ and a single clone of wild type bacteria (OP50) were inoculated in 3 ml LB media overnight at 37 °C and genomic DNA was isolated using Qiagen DNeasy kit (Cat: No. 69504). Purified DNA was sent to Novogene (Novogene Corporation Inc) for sequencing on the Illumina HWI-ST1276 instrument. Data obtained from the Illumina platform was converted into sequencing reads by base calling. The raw data was then filtered to remove low quality reads and adapters. Following the removal of low-quality reads, the FASTQ-files were aligned to the reference genome of E. Coli (MG1655) using BWA aligner ^54,55^. By using SAMtools^56^, the output bam files were sorted and duplicate reads were removed. Afterwards, the index file was created from the bam files. For variant calling, the final bam files were merged and converted into an mpileup file. A variant calling pipeline VarScan.v2, was used for the detection of single nucleotide variants (SNV) and insertions-deletions (indels)^57^. We identified variants in the evolved bacteria (OP50^PQE^) by comparing with genome of the wild type bacteria (OP50) (**Supplementary data 8**).

### Microscopy

Worms were placed in 10% sodium azide (NaN3, vendor) on 3% agarose pads to paralyze them and covered with a glass coverslip. A Leica M165 FC apochromatic corrected fluorescent stereo microscope was used with a PLANAPO 1.0x objective (Leica 10450028) and ET-GFP (EX 470/40, EM 525/50) filter. Images of worms were taken at 6.3x magnification, and with GFP exposure times determined for each reporter strain. Images were taken with the LasX software and analyzed in FIJI for mean fluorescent intensity. Statistics were performed in Graphpad PRISM.

### Inductively coupled plasma mass spectrometry

A single colony of bacteria was inoculated overnight (16 hours) in 50 ml of LB media. Equal O.D of cells from OP50 and OP50^PQE^ was centrifuged at 4400 rpm for 15 minutes. The harvested pellets were washed once with will Milli-Q water, then three times with 1 mM EDTA, followed by one wash with Milli-Q water^58^. Total cellular iron content was then measured as described previously^59^. Briefly, bacterial pellets were treated with 2 mL/g total wet weight nitric acid (Trace metal grade; Fisher) for 24 h, and then digested with 1 mL/g total wet weight hydrogen peroxide (Trace metal grade; Fisher) for 24 h at room temperature. The samples were preserved at 4 °C until quantification of metals. Ultrapure water (VWR Chemicals ARISTAR^®^ULTRA) was used for final sample dilution. Samples were then analyzed using inductively coupled plasma mass spectrometry (ICP-MS) (Perkin Elmer Nexion 2000) using 50 ppb Bismuth as internal standard.

### Statistical analyses

Log-rank test was used to derive p-value for lifespan and paraquat survival assays via OASIS 2 ^60^. On all box plots, the median is shown by the center line, while the upper boundary represents the 75% interquartile range, and the lower boundary represents the 25% interquartile range. GraphPad Prism was used to conduct student t-tests and ANOVAs for two and multiple groups, respectively.

## Supplementary figures

**Fig. S1.**
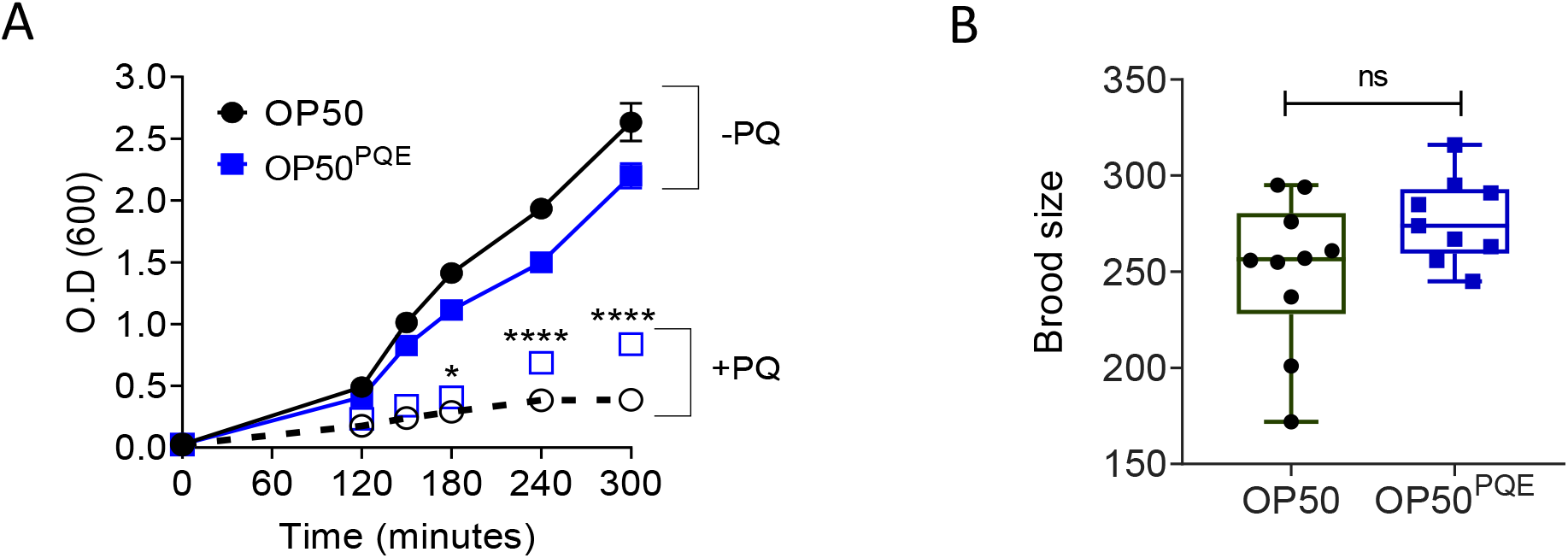
Evolved bacteria resists paraquat and improves worm health. A. Growth curve of wild type bacteria (OP50) and paraquat-evolved bacteria (OP50^PQE^) in the presence and absence of 2.5 mM paraquat (PQ). Growth of OP50^PQE^ is comparable to wild type in normal conditions. Sold line and dotted lines represents growth of bacteria in absence and presence of paraquat, respectively. * and **** represents p-value <0.05 and 0.0001 respectively, calculated using 2-way ANOVA. n = 3 biological replicates, and error bar = mean +/- SD B. Total brood size of worms fed either OP50^PQE^ or OP50 bacteria. ns represents p-value not significant (> 0.05) calculated using unpaired, two-tailed t-test. n = 10 independent worms.

**Fig S2.**
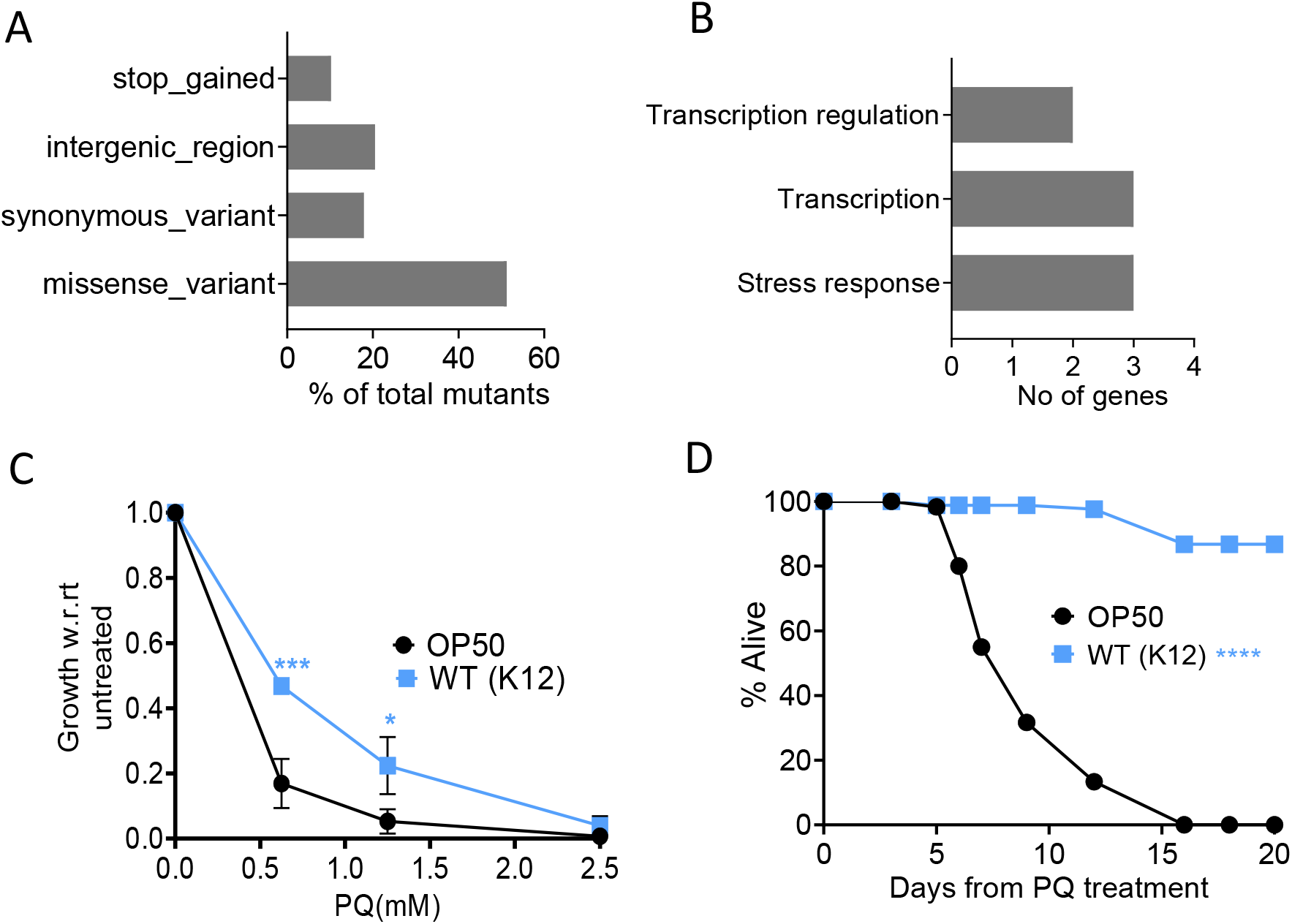
Genomic analysis of evolved bacteria (OP50^PQE^), and comparison of the stress resistance of K12 and OP50 bacteria. A. Bar graph represents the percentage of the different types of mutations present in the OP50^PQE^ bacteria calculated using whole genome sequencing of OP50 and OP50^PQE^ bacteria. B. Biological category for genes mutated in OP50^PQE^ bacteria. Bar graph represents number of genes in different biological terms. C. Dose-dependent effect of different concentrations of paraquat on the growth of OP50 and K12 strain. Error bar represents mean +/- SEM. * and *** represents p-value < 0.05 and < 0.001, respectively, using two-way ANOVA. n = 3 independent biological replicates. D. Survival curve of worms fed OP50 or K12 bacteria in presence of paraquat. Worms fed either of the bacteria were grown till L4 stage and transferred to FUdR plates containing 2.5 mM paraquat. **** represents p-value < 0.0001 calculated using log-rank test.

**Fig S3.**
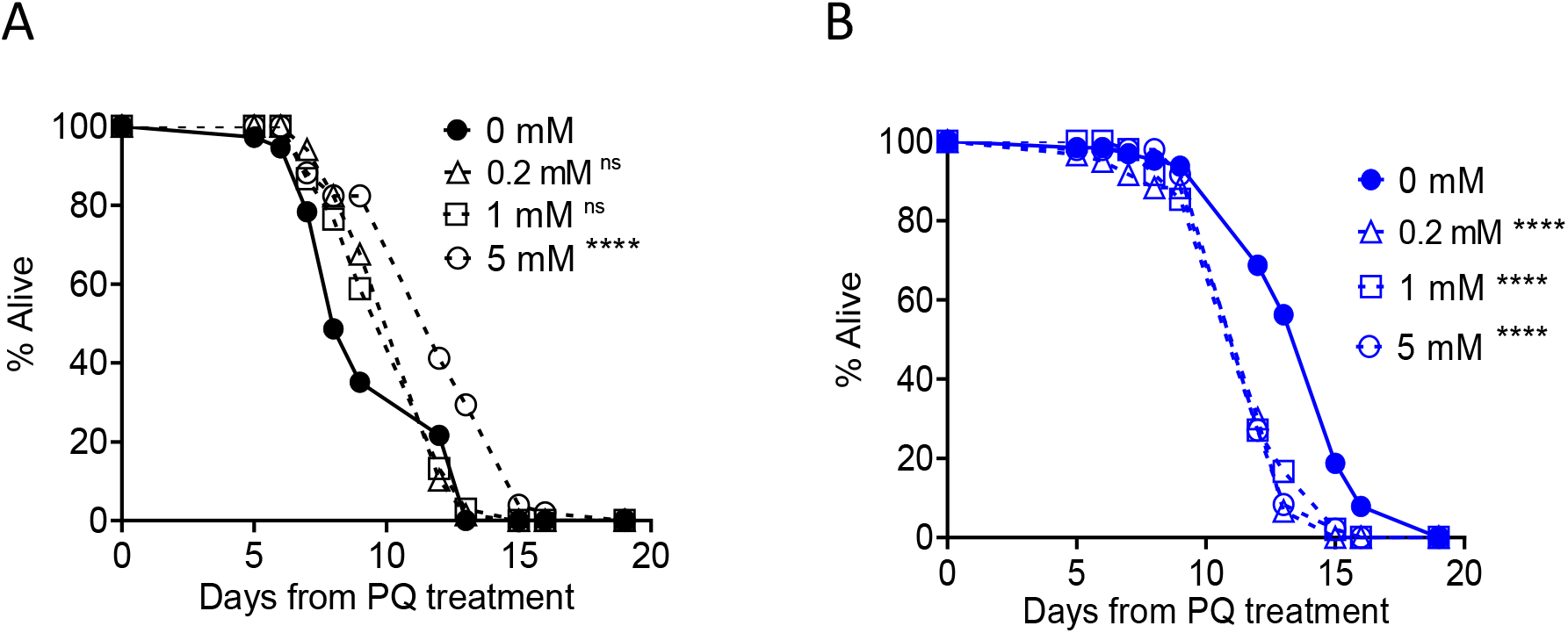
Effect of dietary iron on worm’s resistance to paraquat grown on OP50 vs OP50^PQE^ bacteria. Worms fed OP50 (A) or OP50^PQE^ (B) were transferred to plates containing 2.5 mM paraquat seeded either of the bacteria mixed with different concentrations of iron. **** represent p < 0.0001 respectively calculated using log-rank test.

**Fig. S4.**
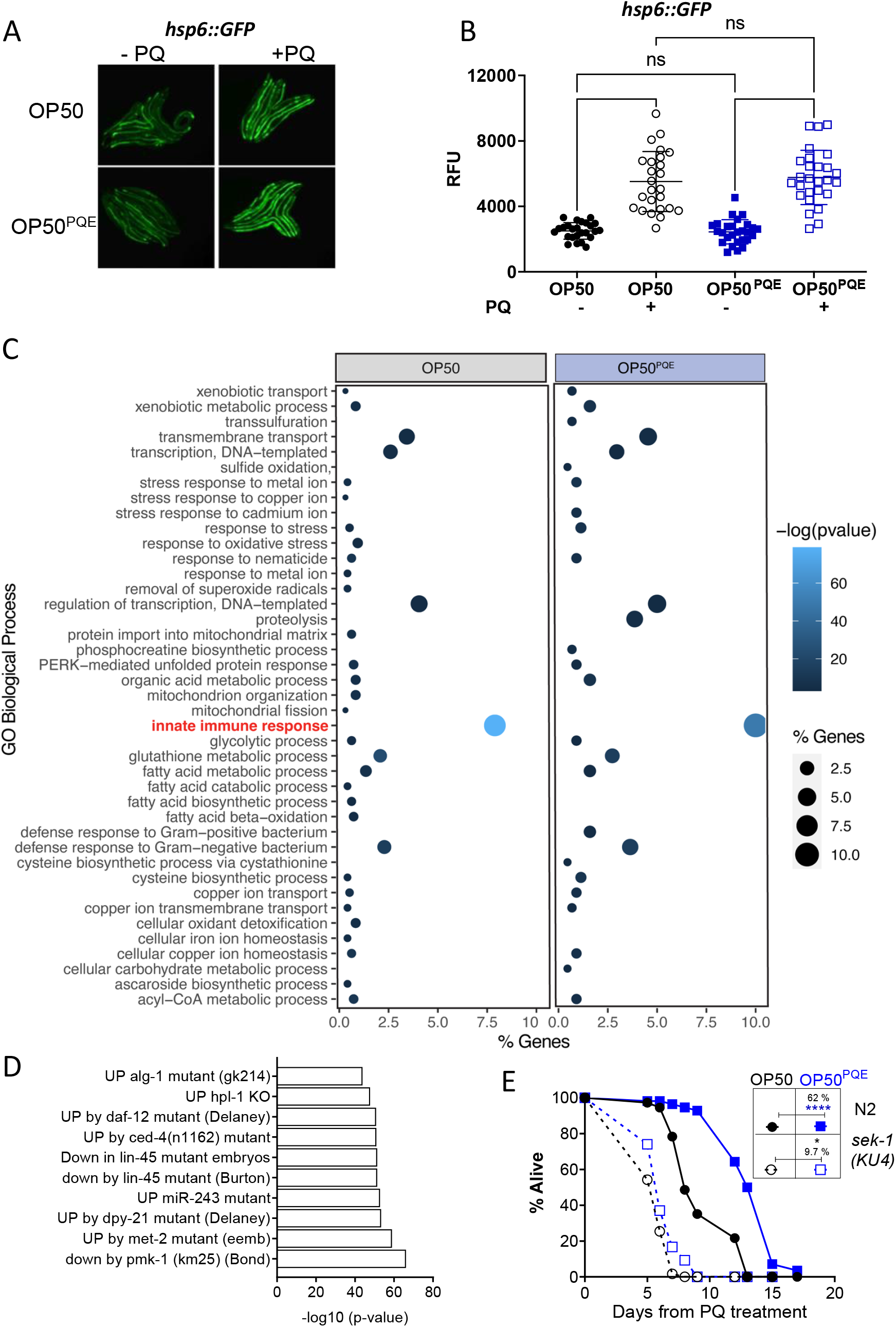
Innate immunity genes are key for differential response of worms fed OP50 or OP50^PQE^ to paraquat exposure. A. Representative images of *hsp-6::GFP* reporter worms grown on either OP50 or OP50^PQE^ with and without exposure to 2.5 mM paraquat. B. Quantification of the fluorescence of *hsp-6p::GFP* reporter worms from F. Error bars represent mean +/- SEM. ns denotes p > 0.05 by an unpaired t-test. C. GO term analysis of RNA-seq results comparing response of worms fed OP50 or OP50^PQE^ to paraquat exposure. Analysis was generated using the DAVID bioinformatics resource (LHRI). D. WormExp analysis highlighting top 10 highly significant datasets that share the genes enriched in worms fed OP50^PQE.^ E. Survival curve of N2 and *sek-1* (KU4) worms fed either OP50 or OP50^PQE^ exposed to 2.5 mM paraquat at the L4 stage. * and **** represents p <0.05 and p < 0.0001, respectively for a log-rank test.

## Supplementary data legends

Supplementary data 1: Stress assay and lifespan analysis

Supplementary data 2: List of high confidence mutations present in OP50^PQE^ bacteria.

Supplementary data 3: Biological terms enriched from genes upregulated due to paraquat in worms fed OP50 or OP50^PQE^ bacteria.

Supplementary data 4: Differentially expressed genes due to paraquat treatment in worms fed OP50 bacteria.

Supplementary data 5: Differentially expressed genes due to paraquat treatment in worms fed OP50^PQE^ bacteria.

Supplementary data 6: Biological terms enriched from genes upregulated in worms fed OP50^PQE^ bacteria.

Supplementary data 7: Differentially expressed genes in worms fed OP50^PQE^ vs OP50 bacteria.

Supplementary data 8: Raw data containing information from the .vcf file generated by genome sequencing of OP50^PQE^ and OP50 bacteria.

## References

1. Kourtis, N., and Tavernarakis, N. (2011). Cellular stress response pathways and ageing: intricate molecular relationships. EMBO J 30, 2520–2531. 10.1038/emboj.2011.162.

2. Li, Z., Zhang, Z., Ren, Y., Wang, Y., Fang, J., Yue, H., Ma, S., and Guan, F. (2021). Aging and age-related diseases: from mechanisms to therapeutic strategies. Biogerontology 22, 165–187. 10.1007/s10522-021-09910-5.

3. Kirkwood, T.B., and Austad, S.N. (2000). Why do we age? Nature 408, 233–238. 10.1038/35041682.

4. Liguori, I., Russo, G., Curcio, F., Bulli, G., Aran, L., Della-Morte, D., Gargiulo, G., Testa, G., Cacciatore, F., Bonaduce, D., and Abete, P. (2018). Oxidative stress, aging, and diseases. Clin Interv Aging 13, 757–772. 10.2147/CIA.S158513.

5. Schieber, M., and Chandel, N.S. (2014). ROS function in redox signaling and oxidative stress. Curr Biol 24, R453–462. 10.1016/j.cub.2014.03.034.

6. Scandalios, J.G. (2002). Oxidative stress responses--what have genome-scale studies taught us? Genome Biol 3, REVIEWS1019. 10.1186/gb-2002-3-7-reviews1019.

7. Oliphant, K., and Allen-Vercoe, E. (2019). Macronutrient metabolism by the human gut microbiome: major fermentation by-products and their impact on host health. Microbiome 7, 91. 10.1186/s40168-019-0704-8.

8. Zheng, D., Liwinski, T., and Elinav, E. (2020). Interaction between microbiota and immunity in health and disease. Cell Res 30, 492–506. 10.1038/s41422-020-0332-7.

9. Pryor, R., Martinez-Martinez, D., Quintaneiro, L., and Cabreiro, F. (2020). The Role of the Microbiome in Drug Response. Annu Rev Pharmacol Toxicol 60, 417–435. 10.1146/annurev-pharmtox-010919-023612.

10. Zhou, Y., Hu, G., and Wang, M.C. (2021). Host and microbiota metabolic signals in aging and longevity. Nat Chem Biol 17, 1027–1036. 10.1038/s41589-021-00837-z.

11. Ghosh, T.S., Shanahan, F., and O’Toole, P.W. (2022). The gut microbiome as a modulator of healthy ageing. Nat Rev Gastroenterol Hepatol 19, 565–584. 10.1038/s41575-022-00605-x.

12. Badal, V.D., Vaccariello, E.D., Murray, E.R., Yu, K.E., Knight, R., Jeste, D.V., and Nguyen, T.T. (2020). The Gut Microbiome, Aging, and Longevity: A Systematic Review. Nutrients 12. 10.3390/nu12123759.

13. Elena, S.F., and Lenski, R.E. (2003). Evolution experiments with microorganisms: the dynamics and genetic bases of adaptation. Nat Rev Genet 4, 457–469. 10.1038/nrg1088.

14. Henry, L.P., Bruijning, M., Forsberg, S.K.G., and Ayroles, J.F. (2021). The microbiome extends host evolutionary potential. Nat Commun 12, 5141. 10.1038/s41467-021-25315-x.

15. Yilmaz, B., and Li, H. (2018). Gut Microbiota and Iron: The Crucial Actors in Health and Disease. Pharmaceuticals (Basel) 11. 10.3390/ph11040098.

16. MacKenzie, E.L., Iwasaki, K., and Tsuji, Y. (2008). Intracellular iron transport and storage: from molecular mechanisms to health implications. Antioxid Redox Signal 10, 997–1030. 10.1089/ars.2007.1893.

17. Anand, N., Holcom, A., Broussalian, M., Schmidt, M., Chinta, S.J., Lithgow, G.J., Andersen, J.K., and Chamoli, M. (2020). Dysregulated iron metabolism in C. elegans catp-6/ATP13A2 mutant impairs mitochondrial function. Neurobiol Dis 139, 104786. 10.1016/j.nbd.2020.104786.

18. Nielsen, B.S., Borregaard, N., Bundgaard, J.R., Timshel, S., Sehested, M., and Kjeldsen, L. (1996). Induction of NGAL synthesis in epithelial cells of human colorectal neoplasia and inflammatory bowel diseases. Gut 38, 414–420. 10.1136/gut.38.3.414.

19. Ng, K.M., Aranda-Díaz, A., Tropini, C., Frankel, M.R., Van Treuren, W., O’Loughlin, C.T., Merrill, B.D., Yu, F.B., Pruss, K.M., Oliveira, R.A., et al. (2019). Recovery of the Gut Microbiota after Antibiotics Depends on Host Diet, Community Context, and Environmental Reservoirs. Cell Host Microbe 26, 650–665.e654. 10.1016/j.chom.2019.10.011.

20. Barreto, H.C., Abreu, B., and Gordo, I. (2022). Fluctuating selection on bacterial iron regulation in the mammalian gut. Curr Biol 32, 3261–3275.e3264. 10.1016/j.cub.2022.06.017.

21. Schwartz, C.J., Giel, J.L., Patschkowski, T., Luther, C., Ruzicka, F.J., Beinert, H., and Kiley, P.J. (2001). IscR, an Fe-S cluster-containing transcription factor, represses expression of Escherichia coli genes encoding Fe-S cluster assembly proteins. Proc Natl Acad Sci U S A 98, 14895–14900. 10.1073/pnas.251550898.

22. Barreto, H.C., Sousa, A., and Gordo, I. (2020). The Landscape of Adaptive Evolution of a Gut Commensal Bacteria in Aging Mice. Curr Biol 30, 1102–1109.e1105. 10.1016/j.cub.2020.01.037.

23. Rajan, M., Anderson, C.P., Rindler, P.M., Romney, S.J., Ferreira Dos Santos, M.C., Gertz, J., and Leibold, E.A. (2019). NHR-14 loss of function couples intestinal iron uptake with innate immunity in. Elife 8. 10.7554/eLife.44674.

24. Coulthard, L.R., White, D.E., Jones, D.L., McDermott, M.F., and Burchill, S.A. (2009). p38(MAPK): stress responses from molecular mechanisms to therapeutics. Trends Mol Med 15, 369–379. 10.1016/j.molmed.2009.06.005.

25. Lee, S., Rauch, J., and Kolch, W. (2020). Targeting MAPK Signaling in Cancer: Mechanisms of Drug Resistance and Sensitivity. Int J Mol Sci 21. 10.3390/ijms21031102.

26. Kim, D.H., Feinbaum, R., Alloing, G., Emerson, F.E., Garsin, D.A., Inoue, H., Tanaka-Hino, M., Hisamoto, N., Matsumoto, K., Tan, M.W., and Ausubel, F.M. (2002). A conserved p38 MAP kinase pathway in Caenorhabditis elegans innate immunity. Science 297, 623–626. 10.1126/science.1073759.

27. Heintz, C., and Mair, W. (2014). You are what you host: microbiome modulation of the aging process. Cell 156, 408–411. 10.1016/j.cell.2014.01.025.

28. Portal-Celhay, C., Bradley, E.R., and Blaser, M.J. (2012). Control of intestinal bacterial proliferation in regulation of lifespan in Caenorhabditis elegans. BMC Microbiol 12, 49. 10.1186/1471-2180-12-49.

29. Zhang, J., Holdorf, A.D., and Walhout, A.J. (2017). C. elegans and its bacterial diet as a model for systems-level understanding of host-microbiota interactions. Curr Opin Biotechnol 46, 74–80. 10.1016/j.copbio.2017.01.008.

30. Gusarov, I., Gautier, L., Smolentseva, O., Shamovsky, I., Eremina, S., Mironov, A., and Nudler, E. (2013). Bacterial nitric oxide extends the lifespan of C. elegans. Cell 152, 818–830. 10.1016/j.cell.2012.12.043.

31. Revtovich, A.V., Lee, R., and Kirienko, N.V. (2019). Interplay between mitochondria and diet mediates pathogen and stress resistance in Caenorhabditis elegans. PLoS Genet 15, e1008011. 10.1371/journal.pgen.1008011.

32. Han, B., Sivaramakrishnan, P., Lin, C.J., Neve, I.A.A., He, J., Tay, L.W.R., Sowa, J.N., Sizovs, A., Du, G., Wang, J., et al. (2017). Microbial Genetic Composition Tunes Host Longevity. Cell 169, 1249–1262.e1213. 10.1016/j.cell.2017.05.036.

33. Choi, H.S., Bhat, A., Howington, M.B., Schaller, M.L., Cox, R.L., Huang, S., Beydoun, S., Miller, H.A., Tuckowski, A.M., Mecano, J., et al. (2023). FMO rewires metabolism to promote longevity through tryptophan and one carbon metabolism in C. elegans. Nat Commun 14, 562. 10.1038/s41467-023-36181-0.

34. Huang, S., Howington, M.B., Dobry, C.J., Evans, C.R., and Leiser, S.F. (2021). Flavin-Containing Monooxygenases Are Conserved Regulators of Stress Resistance and Metabolism. Front Cell Dev Biol 9, 630188. 10.3389/fcell.2021.630188.

35. Dragosits, M., and Mattanovich, D. (2013). Adaptive laboratory evolution -- principles and applications for biotechnology. Microb Cell Fact 12, 64. 10.1186/1475-2859-12-64.

36. Zhou, K.I., Pincus, Z., and Slack, F.J. (2011). Longevity and stress in Caenorhabditis elegans. Aging (Albany NY) 3, 733–753. 10.18632/aging.100367.

37. Partridge, L., Gems, D., and Withers, D.J. (2005). Sex and death: what is the connection? Cell 120, 461–472. 10.1016/j.cell.2005.01.026.

38. Yang, L., Mih, N., Anand, A., Park, J.H., Tan, J., Yurkovich, J.T., Monk, J.M., Lloyd, C.J., Sandberg, T.E., Seo, S.W., et al. (2019). Cellular responses to reactive oxygen species are predicted from molecular mechanisms. Proc Natl Acad Sci U S A 116, 14368–14373. 10.1073/pnas.1905039116.

39. Ding, H., Clark, R.J., and Ding, B. (2004). IscA mediates iron delivery for assembly of iron-sulfur clusters in IscU under the limited accessible free iron conditions. J Biol Chem 279, 37499–37504. 10.1074/jbc.M404533200.

40. Yang, W., Dierking, K., and Schulenburg, H. (2016). WormExp: a web-based application for a Caenorhabditis elegans-specific gene expression enrichment analysis. Bioinformatics 32, 943–945. 10.1093/bioinformatics/btv667.

41. Sheng, Y., Yang, G., Casey, K., Curry, S., Oliver, M., Han, S.M., Leeuwenburgh, C., and Xiao, R. (2021). A novel role of the mitochondrial iron-sulfur cluster assembly protein ISCU-1/ISCU in longevity and stress response. Geroscience 43, 691–707. 10.1007/s11357-021-00327-z.

42. Paul, B.T., Manz, D.H., Torti, F.M., and Torti, S.V. (2017). Mitochondria and Iron: current questions. Expert Rev Hematol 10, 65–79. 10.1080/17474086.2016.1268047.

43. Lill, R., and Mühlenhoff, U. (2006). Iron-sulfur protein biogenesis in eukaryotes: components and mechanisms. Annu Rev Cell Dev Biol 22, 457–486. 10.1146/annurev.cellbio.22.010305.104538.

44. Seyoum, Y., Baye, K., and Humblot, C. (2021). Iron homeostasis in host and gut bacteria - a complex interrelationship. Gut Microbes 13, 1–19. 10.1080/19490976.2021.1874855.

45. Ellermann, M., and Arthur, J.C. (2017). Siderophore-mediated iron acquisition and modulation of host-bacterial interactions. Free Radic Biol Med 105, 68–78. 10.1016/j.freeradbiomed.2016.10.489.

46. Winterbourn, C.C. (1995). Toxicity of iron and hydrogen peroxide: the Fenton reaction. Toxicol Lett 82-83, 969–974. 10.1016/0378-4274(95)03532-x.

47. Zhang, J., Li, X., Olmedo, M., Holdorf, A.D., Shang, Y., Artal-Sanz, M., Yilmaz, L.S., and Walhout, A.J.M. (2019). A Delicate Balance between Bacterial Iron and Reactive Oxygen Species Supports Optimal C. elegans Development. Cell Host Microbe 26, 400–411.e403. 10.1016/j.chom.2019.07.010.

48. Hare, D., Ayton, S., Bush, A., and Lei, P. (2013). A delicate balance: Iron metabolism and diseases of the brain. Front Aging Neurosci 5, 34. 10.3389/fnagi.2013.00034.

49. Le Bourg, E. (2009). Hormesis, aging and longevity. Biochim Biophys Acta 1790, 1030–1039. 10.1016/j.bbagen.2009.01.004.

50. Campos, J.C., Wu, Z., Rudich, P.D., Soo, S.K., Mistry, M., Ferreira, J.C., Blackwell, T.K., and Van Raamsdonk, J.M. (2021). Mild mitochondrial impairment enhances innate immunity and longevity through ATFS-1 and p38 signaling. EMBO Rep 22, e52964. 10.15252/embr.202152964.

51. Miller, H.A., Huang, S., Dean, E.S., Schaller, M.L., Tuckowski, A.M., Munneke, A.S., Beydoun, S., Pletcher, S.D., and Leiser, S.F. (2022). Serotonin and dopamine modulate aging in response to food odor and availability. Nat Commun 13, 3271. 10.1038/s41467-022-30869-5.

52. Parkhomchuk, D., Amstislavskiy, V., Soldatov, A., and Ogryzko, V. (2009). Use of high throughput sequencing to observe genome dynamics at a single cell level. Proc Natl Acad Sci U S A 106, 20830–20835. 10.1073/pnas.0906681106.

53. A Zhang, Y.S., J Li, W Zhang (2019). Improving isobutanol productivity through adaptive laboratory evolution in Saccharomyces cerevisiae. doi.org/ 10.21203/rs.2.19485/v1.

54. Abuín, J.M., Pichel, J.C., Pena, T.F., and Amigo, J. (2015). BigBWA: approaching the Burrows-Wheeler aligner to Big Data technologies. Bioinformatics 31, 4003–4005. 10.1093/bioinformatics/btv506.

55. Li, H., and Durbin, R. (2009). Fast and accurate short read alignment with Burrows-Wheeler transform. Bioinformatics 25, 1754–1760. 10.1093/bioinformatics/btp324.

56. Li, H., Handsaker, B., Wysoker, A., Fennell, T., Ruan, J., Homer, N., Marth, G., Abecasis, G., Durbin, R., and Subgroup, G.P.D.P. (2009). The Sequence Alignment/Map format and SAMtools. Bioinformatics 25, 2078–2079. 10.1093/bioinformatics/btp352.

57. Koboldt, D.C., Chen, K., Wylie, T., Larson, D.E., McLellan, M.D., Mardis, E.R., Weinstock, G.M., Wilson, R.K., and Ding, L. (2009). VarScan: variant detection in massively parallel sequencing of individual and pooled samples. Bioinformatics 25, 2283–2285. 10.1093/bioinformatics/btp373.

58. Huynh, U., Qiao, M., King, J., Trinh, B., Valdez, J., Haq, M., and Zastrow, M.L. (2022). Differential Effects of Transition Metals on Growth and Metal Uptake for Two Distinct. Microbiol Spectr 10, e0100621. 10.1128/spectrum.01006-21.

59. Kremer, D.M., Nelson, B.S., Lin, L., Yarosz, E.L., Halbrook, C.J., Kerk, S.A., Sajjakulnukit, P., Myers, A., Thurston, G., Hou, S.W., et al. (2021). GOT1 inhibition promotes pancreatic cancer cell death by ferroptosis. Nat Commun 12, 4860. 10.1038/s41467-021-24859-2.

60. Han, S.K., Lee, D., Lee, H., Kim, D., Son, H.G., Yang, J.S., Lee, S.V., and Kim, S. (2016). OASIS 2: online application for survival analysis 2 with features for the analysis of maximal lifespan and healthspan in aging research. Oncotarget 7, 56147–56152. 10.18632/oncotarget.11269.

